# Nucleoside-modified mRNA-based influenza vaccines circumvent problems associated with H3N2 vaccine strain egg-adaptation

**DOI:** 10.1101/2022.07.06.499081

**Authors:** Sigrid Gouma, Kaela Parkhouse, Madison Weirick, Hiromi Muramatsu, Norbert Pardi, Steven H.Y. Fan, Drew Weissman, Scott E. Hensley

## Abstract

Most human influenza vaccine antigens are produced in fertilized chicken eggs. Recent H3N2 egg-based vaccine antigens have limited effectiveness, partially due to egg-adaptive substitutions that alter the antigenicity of the hemagglutinin (HA) protein. The nucleoside-modified messenger RNA encapsulated in lipid nanoparticle (mRNA-LNP) vaccine platform is a promising alternative for egg-based influenza vaccines because mRNA-LNP-derived antigens are not subject to adaptive pressures that arise during the production of antigens in chicken eggs. Here, we compared H3N2-specific antibody responses in mice vaccinated with either 3c.2A H3-encoded mRNA-LNP or a conventional egg-based Fluzone vaccine (which included an egg-adapted 3c.2A antigen) supplemented with an MF59-like adjuvant. We tested mRNA-LNP encoding wild-type and egg-adapted 3c.2A H3 antigens. We found that mRNA-LNP encoding wild-type 3c.2A H3 elicited antibodies that neutralized the wild-type 3c.2A H3N2 virus more effectively relative to antibodies elicited by mRNA-LNP encoding egg-adapted 3c2.A H3 or the egg-based Fluzone vaccine. mRNA-LNP expressing either wild-type or egg-adapted 3c2.A H3 protected mice against infection with the wild-type 3c2.A H3N2, whereas the egg-based Fluzone vaccine did not. We found that both mRNA-LNP vaccines elicited high levels of group 2 HA stalk-reactive antibodies that likely contributed to protection *in vivo*. Our studies indicate that nucleoside-modified mRNA-LNP-based vaccines can circumvent problems associated with egg-adaptations with recent 3c2.A H3N2 viruses.

**Summary:** This study shows that the nucleoside-modified messenger RNA encapsulated in lipid nanoparticle (mRNA-LNP) vaccine platform is a promising alternative for egg-based influenza vaccines. We show that mRNA-LNP expressing H3 antigens elicit high levels of antibodies in mice and protect against H3N2 influenza virus infection.

## INTRODUCTION

Influenza vaccine antigens that are prepared in fertilized chicken eggs have limited effectiveness, especially against contemporary H3N2 influenza viruses (1, 2). Recent 3c.2A H3N2 viruses cannot be propagated in eggs without first acquiring adaptive hemagglutinin (HA) substitutions that alter antigenicity. Egg-adapted 3c.2A H3N2 vaccine strains possess a T160K HA substitution that abrogates an antigenically important N-linked glycosylation site at position 158-160 in site B of HA (3). Antibodies elicited by egg-adapted 3c.2A H3N2 vaccine strains react poorly to contemporary wild-type H3N2 viruses (3, 4).

Well before the COVID-19 pandemic, our group and others began developing influenza vaccines based on nucleoside-modified messenger RNA encapsulated in lipid nanoparticles (mRNA-LNP) (5–10). mRNA-LNP encoding a H1 HA elicit high levels of antibodies against the HA head and stalk in mice and ferrets (6). mRNA-LNP HA vaccines elicit sustained germinal center reactions (5) that are able to circumvent the inhibitory effects of maternal antibodies (7). mRNA-LNP-based influenza vaccines have several potential advantages over conventional influenza vaccines. mRNA-LNP influenza vaccines can be easily updated to match antigenically drifted influenza virus strains, and these vaccines are not subject to egg or cell-culture adaptations. Unlike egg-based and cell-based influenza vaccines, antigens from mRNA-LNP-based influenza vaccines are produced in cells of the vaccinee and not subject to selective pressures from egg or cell viral propagation.

Here, we used a murine model to directly compare nucleoside-modified mRNA-LNP vaccines expressing wild-type and egg-adapted H3s with a conventional inactivated egg-based influenza vaccine. We produced mRNA-LNP vaccines expressing wild-type and egg-adapted 3c2.A H3, and we compared these vaccines to a human egg-adapted Fluzone vaccine supplemented with an MF59-like adjuvant. We quantified serum antibody levels after vaccination, and we measured viral lung titers after challenging vaccinated mice with a 3c2.A H3N2 virus.

## RESULTS

### mRNA-LNP expressing wild-type HAs elicit neutralizing antibodies of different specificities compared to mRNA-LNP expressing egg-adapted HAs

3c.2A H3N2 viruses cannot replicate in fertilized chicken eggs without first acquiring a T160K HA substitution, and this egg-adaptation affects antigenicity (3). We created mRNA-LNP vaccines expressing the HA of the A/Hong Kong/4801/2014 3c.2A H3N2 virus, which was the H3N2 component of the 2016-2017 and 2017-2018 Northern Hemisphere influenza vaccine. Egg-adapted A/Hong Kong/4801/2014 vaccine strains possess the T160K HA substitution (3). We created mRNA-LNP expressing the wild-type HA or HA with the egg-adaptive T160K substitution (herein referred to as ‘egg-adapted HA’), and we evaluated these vaccines that differed by a single HA amino acid substitution in BALB/c mice.

We immunized BALB/c mice i.m. with 1 μg or 10 μg of each mRNA-LNP, and we collected serum 1 and 3 months after vaccination. Mice immunized with wild-type HA mRNA-LNP produced higher levels of neutralizing antibodies against the wild-type A/Hong Kong/4801/2014 virus compared to mice immunized with egg-adapted HA mRNA-LNP (**Figure 1A-D**). Conversely, mice immunized with egg-adapted HA mRNA-LNP produced higher levels of neutralizing antibodies against the egg-adapted A/Hong Kong/4801/2014 virus compared to mice immunized with wild-type HA mRNA-LNP (**Figure 1E-H**). Antibody levels elicited by 1μg and 10 μg doses were similar for both wild-type HA mRNA-LNP and egg-adapted HA mRNA-LNP. Antibody levels were similar at 1 and 3 months after vaccination with both wild-type HA mRNA-LNP and egg-adapted HA mRNA-LNP.

**Figure 1.**
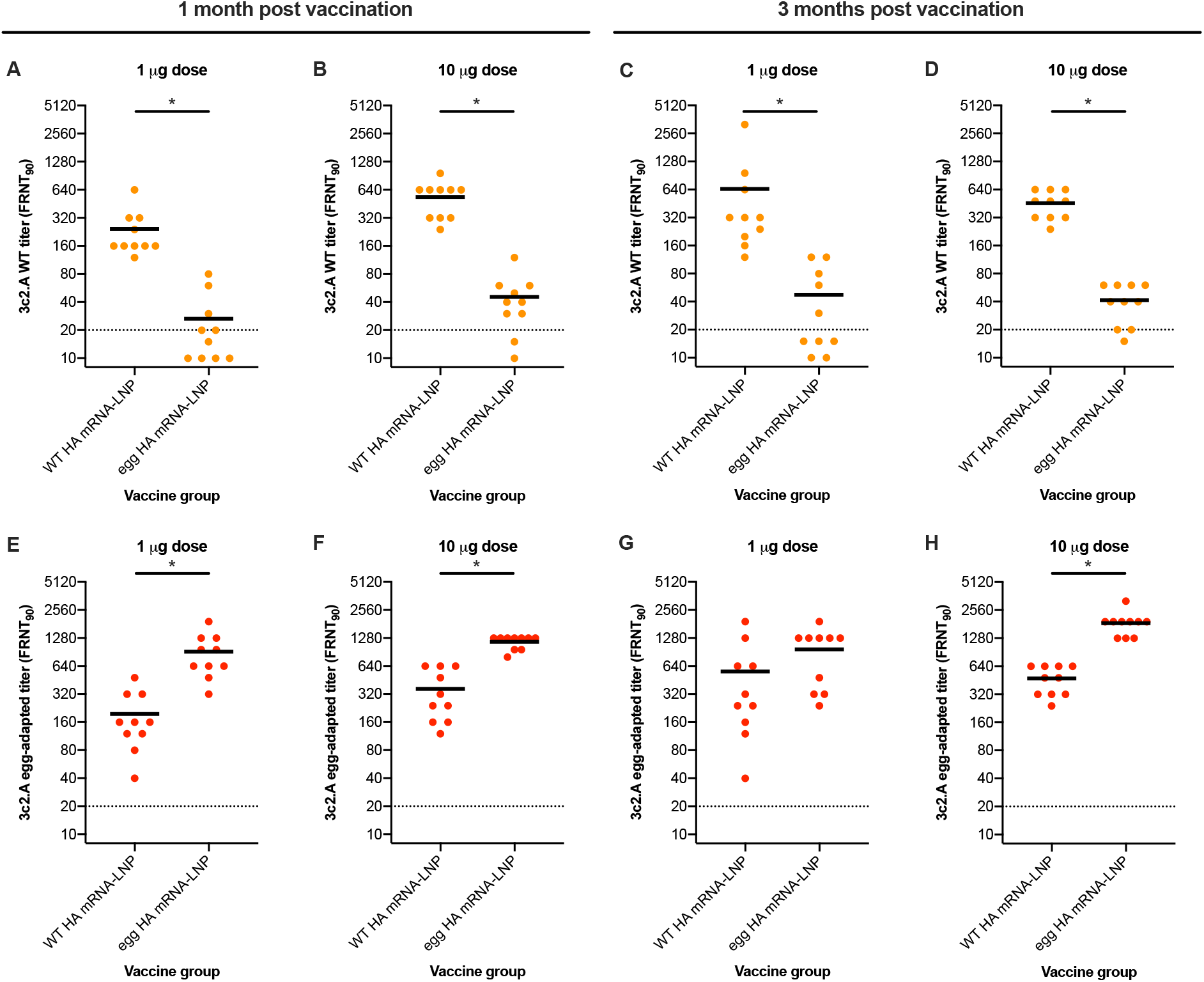
Mice vaccinated with wild-type HA and egg-adapted HA mRNA-LNP produced neutralizing antibodies of different specificities. BALB/c mice were vaccinated with either 1 μg (**A**, **C**, **E**, **G**) or 10 μg (**B**, **D**, **F**, **H**) of each mRNA-LNP vaccine and serum samples were collected 1 and 3 months after vaccination. Neutralizing antibodies titers (FRNT_90_) against wild-type (**A**-**D**) or egg-adapted (**E**-**H**) 3c.2A H3N2 were determined. n=10 mice/group and each symbol represents the mean titer from 2 replicate measurements using serum from 1 animal. Horizontal lines show the mean; dotted line indicates the limit of detection. Statistical analysis: unpaired t test on log_2_ transformed data. * p<0.05.

### Comparison of mRNA-LNP to a conventional egg-based influenza vaccine

We completed additional experiments to directly compare the mRNA-LNP vaccines with the 2017-2018 version of Fluzone. Relative to the wild-type A/Hong Kong/4802/2014 HA, the H3N2 component of the 2017-2018 Fluzone vaccine possesses N96S, T160K, L194P, and D225N egg-adapted HA substitutions. We added the MF59-like adjuvant Addavax to the Fluzone vaccine for our experiments, because conventional inactivated vaccines are poorly immunogenic in mice without adjuvant. We vaccinated BALB/c mice with a total volume of 50 μL Fluzone/Addavax mixture, which contains 0.75 μg of each of the influenza antigens (H1, H3, and 2 influenza B virus HAs). In parallel, we vaccinated additional groups of BALB/c mice with 10 μg of mRNA-LNP expressing wild-type or egg-adapted HAs. We obtained serum 28 days after vaccination for serological analyses.

Similar to our initial vaccine dose titration experiments, mice immunized with wild-type HA mRNA-LNP produced high levels of neutralizing antibodies against the wild-type A/Hong Kong/4801/2014 virus (**Figure 2A**), while mice immunized with egg-adapted HA mRNA-LNP produced high levels of neutralizing antibodies against the egg-adapted A/Hong Kong/4801/2014 virus (**Figure 2B**). Mice immunized with Fluzone + Addavax produced antibodies that could weakly neutralize the egg-adapted A/Hong Kong/4801/2014 virus (**Figure 2B**), and these antibodies had no neutralizing activity against the wild-type A/Hong Kong/4801/2014 virus (**Figure 2A**). The egg-adapted virus in our study possessed only the T160K HA substitution, whereas the Fluzone egg-adapted 3c2.A virus possessed the N96S, L194P, and D225N substitutions in addition to the T160K substitution. For this reason, we repeated neutralization assays using a 3c2.A virus with all 4 HA substitutions that were in the Fluzone egg-adapted 3c.2A virus. Antibodies elicited by Fluzone + Addavax neutralized the Fluzone egg-adapted 3c2.A virus (with all 4 egg-adapted substitutions) better (**Figure 2C**) compared to the egg-adapted 3c2.A virus with only the T160K HA substitution (**Figure 2B**). Antibodies elicited by wild-type HA mRNA-LNP had similar neutralizing activity against the Fluzone egg-adapted 3c2.A virus (**Figure 2C**) and the 3c2.A virus with only the T160K substitution (**Figure 2B**). Antibodies elicited by egg-adapted HA mRNA-LNP also had similar neutralizing activity against the Fluzone egg-adapted 3c2.A virus (**Figure 2C**) and the 3c2.A virus with only the T160K substitution (**Figure 2B**).

**Figure 2.**
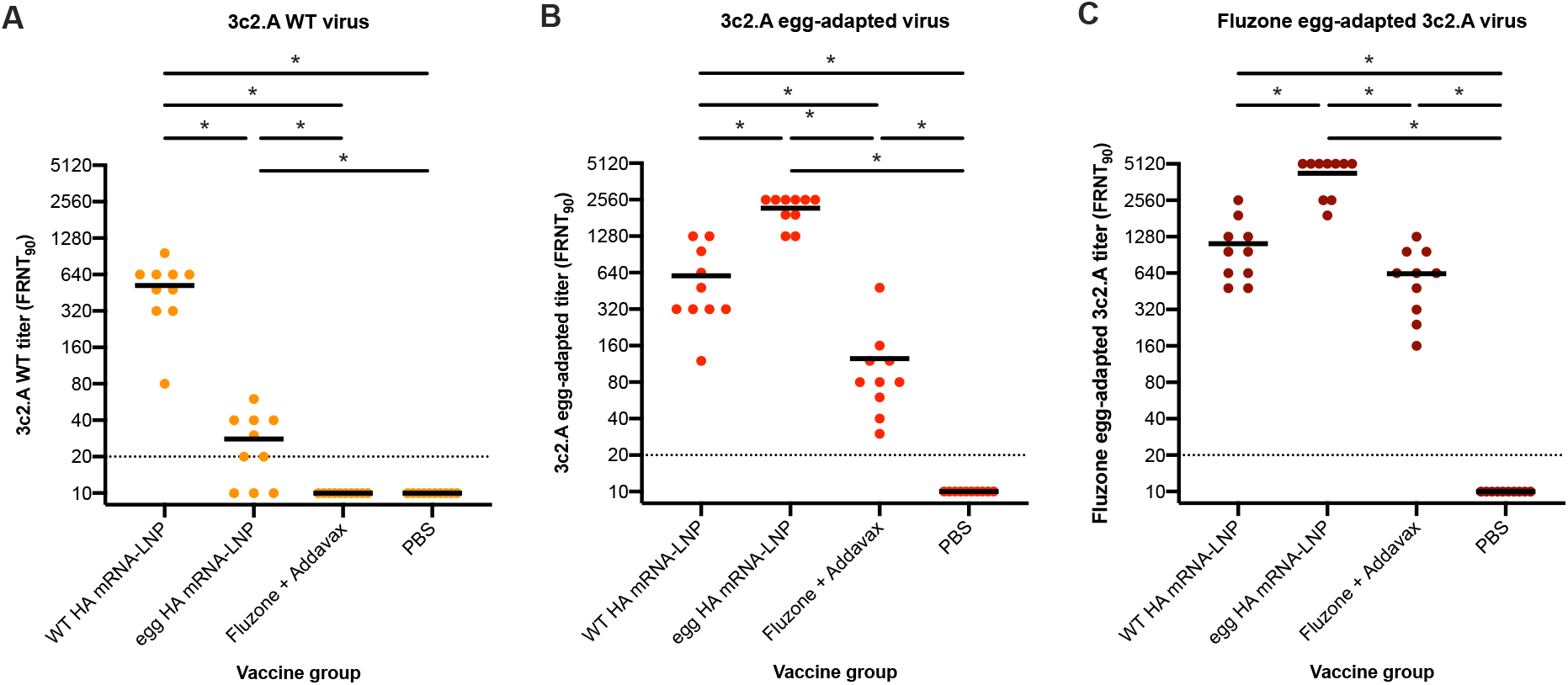
Mice vaccinated with Fluzone + Addavax do not mount neutralizing antibodies against wild type 3c2.A H3N2 viruses. BALB/c mice were vaccinated with either 10 μg of each mRNA-LNP vaccine or 50 μL Fluzone/Addavax mixture, and serum samples were collected 28 days after vaccination. Neutralizing antibodies titers (FRNT_90_) against wild-type (**A**), egg-adapted (with only the T160K HA substitution) (**B**), and Fluzone egg-adapted (with 4 HA substitutions) (**C**) 3c.2A H3N2 viruses were determined. Data from two independent experiments were combined. n=10 mice/group and each symbol represents the mean titer from 2 replicate measurements using serum from 1 animal. Horizontal lines show the mean; dotted line indicates the limit of detection. Statistical analysis: one-way ANOVA with Bonferroni correction on log_2_ transformed data. * p<0.05.

### HA mRNA-LNP vaccines elicit antibodies that target conserved epitopes in H3 stalk

We previously demonstrated that mRNA-LNP vaccines expressing H1 antigens elicit neutralizing antibodies, as well as antibodies that bind to conserved epitopes within H1 (group 1) stalk proteins (6). We completed additional ELISAs with serum collected 28 days after vaccination to determine if the H3 mRNA-LNP vaccines in our study elicit antibodies that bind to conserved epitopes in H3 (group 2) stalk proteins. We quantified antibodies reactive to a wild-type 3c2.A full-length recombinant HA protein (**Figure 3A**) and a headless H3 recombinant protein (**Figure 3B**). The wild-type HA mRNA-LNP vaccine and egg-adapted HA mRNA-LNP vaccine elicited high levels of antibodies that bound to the 3c2.A full-length recombinant HA protein, whereas the Fluzone + Addavax vaccine elicited lower levels of antibodies that bound to this HA (**Figure 3A**). Both mRNA-LNP vaccines elicited similar levels of antibodies that bound to the headless H3 stalk, while the Fluzone + Addavax vaccine did not elicit detectable levels of H3 stalk-binding antibodies (**Figure 3B**).

**Figure 3.**
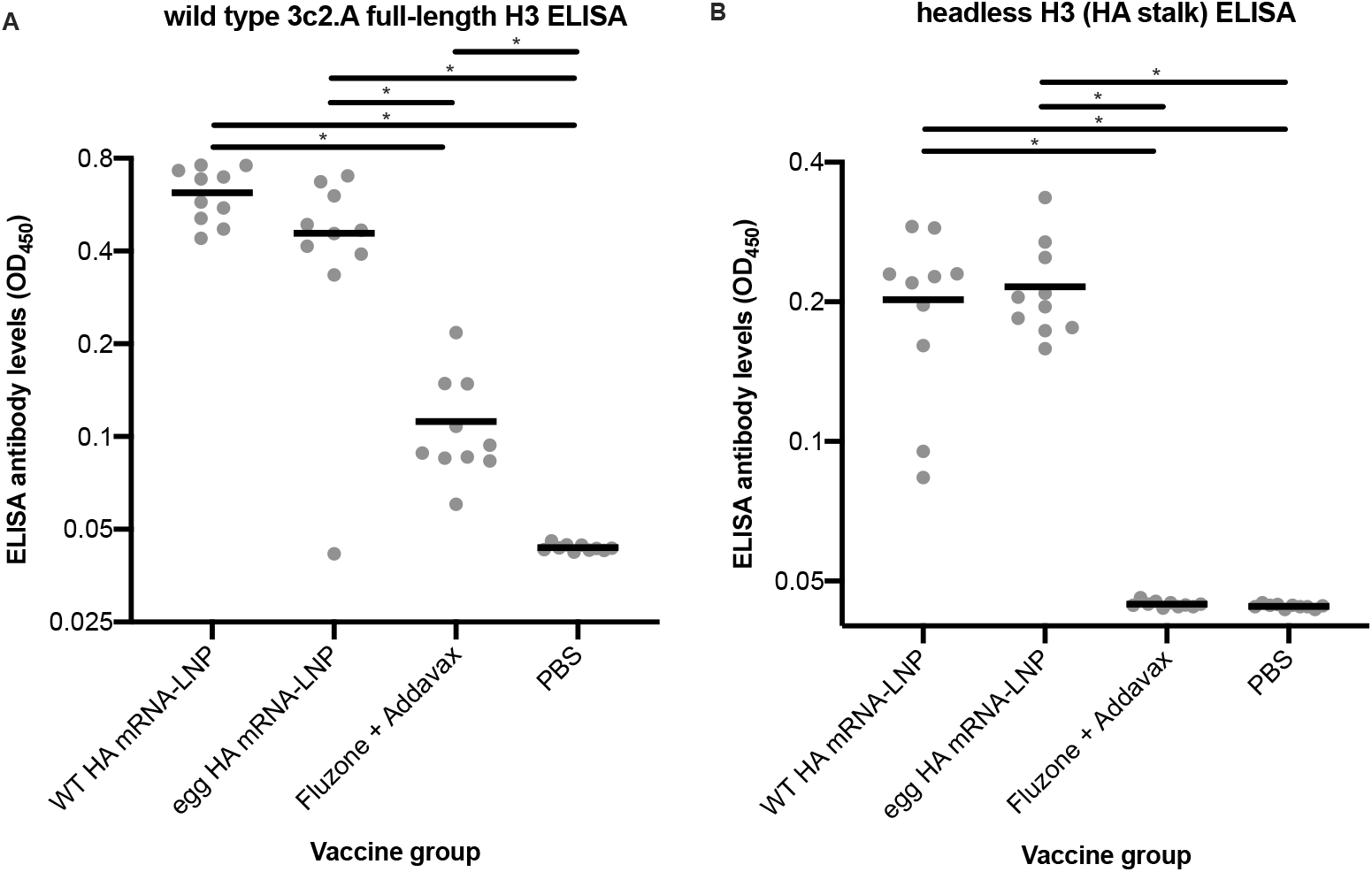
mRNA-LNP vaccines, but not Fluzone + Addavax, elicit H3 stalk-reactive antibodies. BALB/c mice were vaccinated with either 10 μg of each mRNA-LNP vaccine or 50 μL Fluzone/Addavax mixture, and serum samples were collected 28 days after vaccination. ELISA antibody levels reactive to wild-type 3c2.A full-length recombinant HA (**A**) and headless H3 (**B**) were determined. n=10 mice. Horizontal lines show the mean. Statistical analysis: one-way ANOVA with Bonferroni correction. * p<0.05.

### Antibodies elicited by HA mRNA-LNP prevent replication of wild-type 3c2.A H3N2 viruses in mice

We completed experiments to determine if antibodies elicited by HA mRNA-LNP and Fluzone + Addavax vaccines protect mice from infection with a wild-type 3c2.A virus. For this, we passively transferred serum from vaccinated mice into previously unexposed mice, and then we intranasally infected these mice with viruses that possessed the wild-type A/Hong Kong/4801/2014 HA. The challenge virus used in these experiments possessed internal genes from the mouse-adapted A/Puerto Rico/8/1934 strain to facilitate viral replication. Since viruses with the wild-type 3c2.A HA do not cause weight loss and measurable disease in mice, we isolated lung tissue 2 days after infection and quantified lung viral titers as a proxy for influenza virus disease. Viral lung titers were the highest in control mice that received sera from unvaccinated mice and in mice that received sera from mice vaccinated with Fluzone +

Addavax (**Figure 4**). As expected based on our neutralizing antibody data, viral lung titers were significantly lower in mice that received sera from mice vaccinated with the wild-type HA mRNA-LNP vaccine. Viral lung titers were also low in mice that received sera from mice vaccinated with the egg-adapted HA mRNA-LNP vaccine. This was surprising because the egg-adapted HA mRNA-LNP vaccine elicited low levels of neutralizing antibodies against the 3c.2A wild-type virus (**Figure 2A**). Since we found that both mRNA-LNP vaccines elicited similar levels of H3 stalk binding antibodies (**Figure 3B**), these experiments suggest that non-neutralizing antibodies elicited by H3 mRNA-LNP likely contribute to protection when the mRNA-LNP H3 vaccine antigens are mismatched to challenge strains.

**Figure 4.**
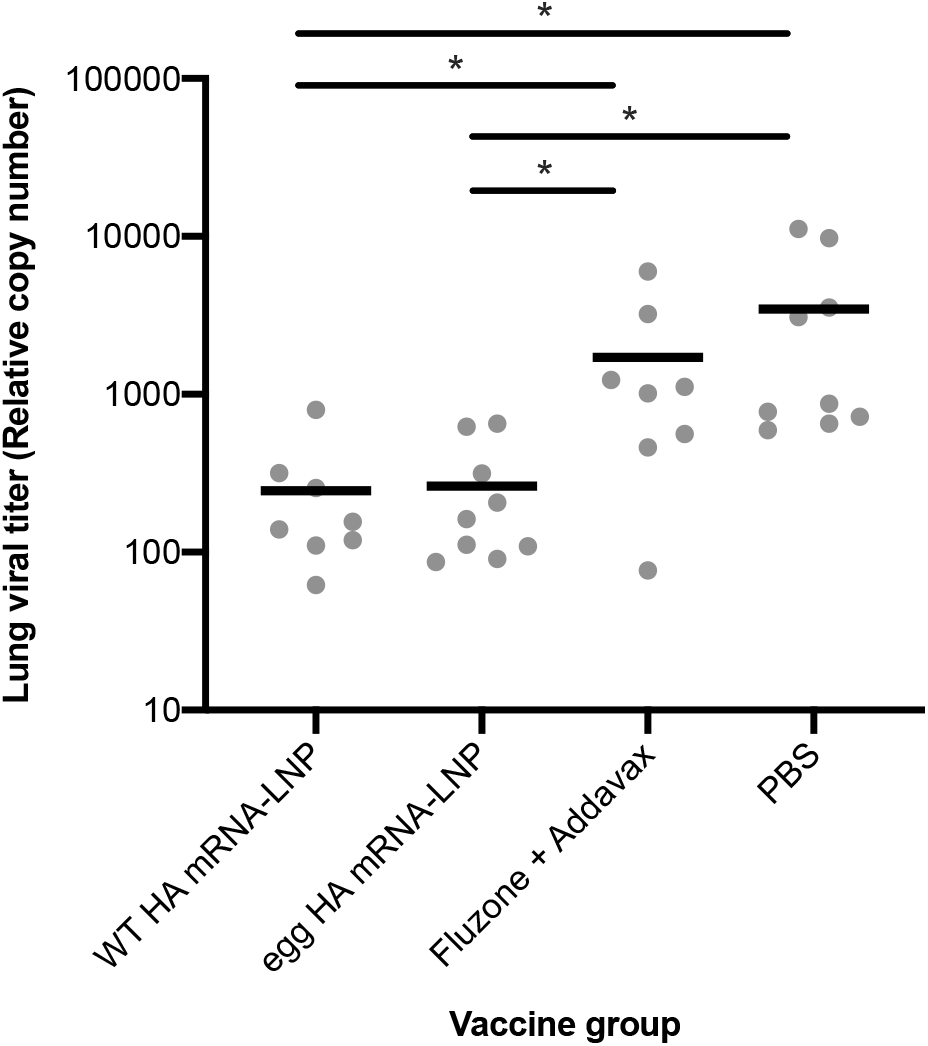
Antibodies elicited by HA mRNA-LNP reduce wild-type 3c2.A H3N2 virus replication in mice. Serum was collected from vaccinated mice 28 days after vaccination and then transferred intraperitoneally to naive unexposed mice 4 hours before i.n. infection with wild-type 3c.2A H3N2 virus. Relative viral copy numbers in lungs were measured by RT-PCR 2 days after viral infections. Data from two independent experiments were combined. n=8-9 mice and each symbol represents the mean titer from 2 replicate measurements from samples from 1 animal. Horizontal lines show the mean. Statistical analysis: one-way ANOVA with Bonferroni correction on log_10_ transformed data. * p<0.05.

## DISCUSSION

Conventional inactivated and subunit influenza vaccines dramatically reduce hospitalizations and deaths caused by influenza viruses, but there is certainly room for improvement. Influenza viruses are constantly changing and vaccines are less effective when vaccine strains are mismatched to circulating strains (2). Vaccine strains are typically chosen ~9 months prior to an influenza season, and it is not always possible to accurately predict which viral strains will dominate. To complicate matters further, influenza virus HA antigens sometimes change during the vaccine manufacturing process. Viruses often acquire adaptive HA mutations that facilitate replication in cells used for vaccine antigen production, and these adaptative mutations can affect antigenicity. Egg-adaptations have been particularly problematic for recent 3c.2A H3N2 viruses, since these viruses cannot be propagated in fertilized chicken eggs without first acquiring a T160K HA substitution that alters antibody binding (3).

Nucleoside-modified mRNA-LNP vaccines have been a useful tool against SARS-CoV-2 and these vaccines show promise for other infectious diseases, including influenza viruses (11). These vaccines can be updated faster than most conventional influenza vaccine approaches, which can potentially limit vaccine strain mismatches. mRNA-LNP vaccine antigens are produced in cells of vaccinees and are therefore not subject to selective pressures related to vaccine production. In this report, we created a nucleoside-modified mRNA-LNP vaccine expressing a wild-type H3 antigen that is difficult to produce in fertilized chicken eggs, and we show that this vaccine is more effective than an adjuvanted conventional influenza vaccine in a murine model.

We also tested an mRNA-LNP vaccine expressing an egg-adapted H3 antigen. While our mRNA-LNP vaccine expressing wild-type H3 antigen elicited higher levels of neutralizing antibodies against wild-type H3N2 relative to our mRNA-LNP vaccine expressing egg-adapted H3 antigen, both vaccines prevented replication of wild-type H3N2 virus in mice. Similar to what we previously found for group 1 H1 viruses (6), we found that H3 mRNA-LNP elicited group 2 HA stalk-reactive antibodies. mRNA-LNP vaccines elicit sustained germinal center reactions (5), which likely promote antibody responses against epitopes that are usually subdominant, such as antibodies directed against the HA stalk. Antibodies that recognize non-neutralizing epitopes of the HA stalk can provide protection through mechanisms that require Fc receptor interactions (12).

Collectively, our studies show that H3 mRNA-LNP vaccines can overcome problems associated with egg-adaptations and are also effective when the HA is antigenically mismatched to challenge strains. Further studies should evaluate different H3 mRNA-LNP vaccines in other animal models and humans. There is an urgent need to speed up the process of updating specific mRNA-LNP antigens for human use. There has been a major delay updating SARS-CoV-2 vaccine antigens and it will be important to accelerate this process for H3 mRNA-LNP vaccines since H3N2 viruses continuously undergo antigenic drift.

## MATERIALS AND METHODS

### Vaccines

mRNA-LNP were produced as previously described (13). mRNA-LNP were diluted in 50 μL of sterile PBS and animals were vaccinated intramuscularly (i.m.). Quadrivalent Fluzone 2017-2018 vaccine (Sanofi Pasteur) was mixed with Addavax (Invivogen), an MF59-like adjuvant, at a 1:1 ratio. Animals were injected with a total volume of 50 μL Addavax/MF59 mixture, which contains 0.75 μg of each of the influenza antigens (H1, H3, and 2 influenza B virus HAs).

### Viruses

Viruses expressing the wild-type HA from A/Hong Kong/4801/2014 (3c2.A), the egg-adapted HA from A/Hong Kong/4801/2014 (with only the T160K substitution), or the Fluzone egg-adapted HA from A/Hong Kong/4801/2014 (with the N96S, T160K, L194P and D225N substitutions) were generated through reverse genetics. The NA from the wild-type A/Hong Kong/4801/2014 strain was included in all 3 reverse-genetics derived viruses. Viruses were transfected in co-cultures of 293T and MDCK-SIAT1 cells with plasmids (pHW2000-or pDZ-based) encoding all 8 influenza virus gene segments. Viruses were rescued using the internal genes from A/Puerto Rico/8/1934 in combination with HA and NA genes of interest. Transfection supernatants were harvested 3 days after transfection and stored at −80°C. HA and NA genes of all viruses were sequenced to confirm that no additional mutations were introduced.

### Animals

BALB/c mice (7-9 week old female) were obtained from Charles River Laboratories. All animal procedures were approved and performed in accordance with the Wistar Institute IACUC guidelines. Vaccines were injected i.m. into the quadriceps muscle. After 28 days, serum was collected by either submandibular bleeding or, if used for passive transfer into naive mice, cardiac bleeding, using serum collection tubes (Sarstedt). For passive transfer experiments, sera from each vaccine group were pooled (n=5) and naive mice were intraperitoneally injected with 150 μL pooled serum. After 4 hours, serum was collected by submandibular bleeding to determine whether antibodies were efficiently transferred, and we excluded mice that had no detectable antibodies upon passive transfer. For some experiments we anesthetized and then intranasally infected mice with 10,000 FFU of H3N2 viruses with the A/Hong Kong/4801/2014 HA diluted in sterile PBS in a total volume of 50 μL. Mice were euthanized 2 days later to harvest and homogenize lungs and determine influenza virus RNA levels.

### Foci Reduction Neutralization Test (FRNT)

RDE-treated serum samples were serially diluted in MEM (Gibco). Viruses were diluted to a concentration of approximately 300 focus-forming units per well and then incubated with serum for 1 hour at room temperature. Confluent monolayers of MDCK-SIAT1 cells were washed with MEM before virus-serum mixtures were added to each well. Cells were incubated for 1 hour at 37°C in 5% CO_2_ and then washed with MEM, after which an overlay of 1.25% Avicel in MEM supplemented with 0.2% gentamicin and 1% 1M HEPES was added to the cells. After an 18 hour incubation at 37°C in 5% CO_2_, cells were fixed with 4% paraformaldehyde and then lysed with 0.5% Triton X-100 in PBS for 7 minutes followed by blocking with 5% milk in PBS. Plates were washed with water before 50 μL of anti-NP monoclonal antibody IC5-1B7 (BEI) was added to each well. After incubation, plates were washed with water and anti-mouse peroxidase-conjugated secondary antibody (Fisher) was added to each well. Plates were incubated, washed and TMB substrate (Kirkegaard & Perry Laboratories) was added for visualization of the foci. Plates were imaged and foci were quantified using an ELISpot reader. FRNT90 titers were reported as the reciprocal of the highest dilution of sera that reduced the number of foci by at least 90%, relative to control wells that had no serum. An anti-A/Hong Kong/4801/2014 “in-house” polyclonal antibody control was included in each assay run.

### Viral load quantification (qRT-PCR)

Mouse lungs were homogenized and RNA was extracted using the Qiagen RNeasy Mini Kit (Qiagen). Extracted RNA was used for cDNA synthesis using Superscript III reverse transcriptase (Invitrogen) and random hexamers (Invitrogen). cDNA was used for amplification using the Taqman Universal PCR Master Mix (Applied Biosystems). Primers and probe were used as described previously (14): Fwd: 5’GGACTGCAGCGTAGACGCTT; Rev: 5’CATCCTGTTGTATATGAGGCCCAT; probe: 5’CTCAGTTATTCTGCTGGTGCACTTGCCA.

The following program was used: 50°C for 2 minutes, 95°C for 10 minutes, followed by 40 cycles of 95°C for 15 seconds and 60°C for 1 minute.

### Full-length H3 enzyme-linked immunosorbent assay (ELISA)

ELISA plates were coated overnight at 4°C with wild-type A/Hong Kong/4801/2014 full-length recombinant HA protein. The next day, ELISA plates were blocked for 2 hours with PBS containing 3% BSA. Plates were washed with PBS containing 0.1% Tween 20 (PBS-T) and 50 μL serum dilution was then added to each well. After 2 hours of incubation, plates were washed with PBS-T and 50 μL horseradish peroxidase (HRP)-conjugated anti-mouse secondary antibody diluted in PBS containing 1% BSA was added to each well. After an hour incubation, plates were washed with PBS-T and SureBlue TMB substrate (KPL) was added to develop the plates. After 5 minutes the developing process was stopped with 250 mM hydrochloric acid. Plates were read at an optical density (OD) of 450 nm using the SpectraMax 190 microplate reader (Molecular Devices).

### Headless H3 ELISA

ELISA plates were coated overnight at 4°C with streptavidin. The next day, plates were washed with PBS-T incubated for an hour with biotinylated headless H3 (15). Plates were washed with PBS-T and then blocked for an hour with TBS supplemented with 0.05% Tween and 1% BSA. Plates were again washed with PBS-T before 50 μL serum dilution was added to each well. After an hour incubation, plates were washed with PBS-T and 50 μL horseradish peroxidase (HRP)-conjugated anti-mouse secondary antibody diluted in TBS containing 0.05% Tween and 0.1% BSA was added to each well. After an hour incubation, plates were washed with PBS-T and SureBlue TMB substrate (KPL) was added to develop the plates. After 5 minutes the developing process was stopped with 250 mM hydrochloric acid. Plates were read at an optical density (OD) of 450 nm using the SpectraMax 190 microplate reader (Molecular Devices).

### Statistical analysis

Data were compared between vaccine groups using one-way ANOVA and two-way ANOVA corrected for multiple comparisons (Bonferroni method). Antibody titers were log_2_-transformed and viral lung titers were log_10_-transformed prior to analysis. Analyses were performed using GraphPad Prism version 7. P values <0.05 were considered as statistically significant.

## ACKNOWLEDGEMENTS

This project has been funded in part with Federal funds from the National Institute of Allergy and Infectious Diseases, National Institutes of Health, Department of Health and Human Services, under Contract No. 75N93021C00015, and Grant No. 1R01AI108686. SEH holds an Investigators in the Pathogenesis of Infectious Disease Awards from the Burroughs Wellcome Fund. We thank Adrian McDermott (NIH) for providing the headless H3 construct.

## COMPETING INTERESTS

SEH reports receiving consulting fees from Sanofi Pasteur, Lumen, Novavax, and Merck. In accordance with the University of Pennsylvania policies and procedures and our ethical obligations as researchers, we report that SEH, N.P. and DW are named on patents that describe the use of nucleoside-modified mRNA as a platform to deliver therapeutic proteins and vaccines. We have disclosed those interests fully to the University of Pennsylvania, and we have in place an approved plan for managing any potential conflicts arising from licensing of our patents. SHYF is an employee of Acuitas Therapeutics, a company focused on the development of lipid nanoparticulate nucleic acid delivery systems for therapeutic applications.

